# Expectations about motion direction affect perception and anticipatory smooth pursuit differently

**DOI:** 10.1101/2020.11.02.365338

**Authors:** Xiuyun Wu, Austin C. Rothwell, Miriam Spering, Anna Montagnini

**Affiliations:** Graduate Program in Neuroscience, University of British Columbia, Vancouver, BC, Canada; Department of Ophthalmology & Visual Sciences, University of British Columbia, Vancouver, BC, Canada; Djavad Mowafaghian Center for Brain Health, University of British Columbia, Vancouver, BC, Canada; Institute for Computing, Information and Cognitive Systems, University of British Columbia, Vancouver, BC, Canada; Institut de Neurosciences de la Timone, CNRS and Aix-Marseille Université, Marseille, France

## Abstract

Smooth pursuit eye movements and visual motion perception rely on the integration of current sensory signals with past experience. Experience shapes our expectation of current visual events and can drive eye movement responses made in anticipation of a target, such as anticipatory pursuit. Previous research revealed consistent effects of expectation on anticipatory pursuit—eye movements follow the expected target direction or speed—and contrasting effects on motion perception, but most studies considered either eye movement or perceptual responses. The current study directly compared effects of direction expectation on perception and anticipatory pursuit within the same direction discrimination task to investigate whether both types of responses are affected similarly or differently. Observers (*n* = 10) viewed high-coherence random-dot kinematograms (RDKs) moving rightward and leftward with a probability of 50, 70, or 90% in a given block of trials to build up an expectation of motion direction. They were asked to judge motion direction of interleaved low-coherence RDKs (0-15%). Perceptual judgements were compared to changes in anticipatory pursuit eye movements as a function of probability. Results show that anticipatory pursuit velocity scaled with probability and followed direction expectation (attraction bias), whereas perceptual judgments were biased opposite to direction expectation (repulsion bias). Control experiments suggest that the repulsion bias in perception was not caused by retinal slip induced by anticipatory pursuit, or by motion adaptation. We conclude that direction expectation can be processed differently for perception and anticipatory pursuit.

## Introduction

How we perceive and interact with the visual world depends not only on current visual input but also on our experience with past sensory events. In Bayesian inference, this experience informs a prior—one’s expectation of the probability of an event before any sensory evidence is present (de Lange et al. 2018; Seriès and Seitz 2013). This study investigates how visual motion priors, based on long-term experience, affect visual perception and movement, and whether these two outcomes are controlled in the same way or differently by expectation. We use smooth pursuit eye movements—the eyes’ continuous response to moving objects—as a model system for visually-guided movement to investigate this question. Smooth pursuit eye movements are closely related to the perception of visual motion (Gegenfurtner 2016; Schütz et al. 2011; Spering and Montagnini 2011). They rely on the integration of current motion information with priors based on experience across just a few trials or across a longer-term context (Darlington et al. 2017; Deravet et al. 2018; Yang et al. 2012). Moreover, smooth pursuit can be triggered by the expectation of a certain motion direction even before the object’s motion onset, a phenomenon known as anticipatory smooth pursuit (Kowler et al. 1984, 2019).

Previous research has revealed highly consistent effects of expectation on pursuit but contrasting effects on motion perception. For example, anticipatory pursuit can be triggered when observers repeatedly view stimuli moving into the same direction (Kowler 1989; Kowler et al. 2019). The eyes are then attracted to the expected motion direction prior to the onset of the stimulus (attraction bias in direction). These responses are not purely habitual but finely tuned to the strength of expectation (Damasse et al. 2018; Jarrett and Barnes 2002; Santos and Kowler 2017). Congruently, anticipatory pursuit velocity is proportional to the average velocity of the target across previous trials (attraction bias in speed), and strongly affected by events in the previous two trials (Maus et al. 2015). Furthermore, Bayesian integration models have been used to describe how priors would lead to attraction effects in visually-guided pursuit when combined with noisy visual motion signals (Behling and Lisberger 2020; Darlington et al. 2017; Deravet et al. 2018).

By contrast, perceptual studies have found evidence for both an attraction bias as well as for responses to be repelled away from the expected direction (repulsion bias; Jazayeri and Movshon 2007). In studies reporting attraction biases, estimates of direction, velocity, or orientation in a current trial are affected by events in previous trials such that an observer’s perception would be biased in line with the motion information observed in previous trials. Such biases can build up quickly, within a few trials (Alais et al. 2017; Cicchini et al. 2018) or can be based on implicitly learning the statistical properties of a stimulus environment over many trials (Chalk et al. 2010; Kok et al. 2013; Seriès and Seitz 2013), as described by Bayesian integration models. Perceptual repulsion biases have been observed across different visual tasks and features. In a speed estimation task in which observers had to judge whether the speed in the current trial was faster or slower than the average speed across all previous trials, observers tended to overestimate a current target’s velocity when the average velocity across previous trials was slow and vice versa for fast velocity (Maus et al. 2015). Similar repulsion biases have been found in studies in which observers had to adjust the orientation of a test stimulus relative to an inducer stimulus, when both stimulus orientations differed by more than 60° between the previous and current trial (Fritsche et al. 2020). In this scenario, observers’ adjustment responses were sometimes repelled away from the previous trial’s stimulus orientation. In sum, expectation built across different timescales can result in a perceptual bias either in the same direction as the cue, prompt, or adaptor (attraction bias), or in the opposite direction (repulsion bias).

The current study directly compared effects of direction expectation on perception and anticipatory pursuit within the same trials to investigate whether both types of responses are affected similarly or differently, and how they interact. On one hand, attraction biases are commonly found in pursuit, and in most perceptual studies that did not use adjustment tasks or reference comparisons. On the other hand, one study that directly compared velocity expectation effects reported opposite biases in speed discrimination and anticipatory pursuit (Maus et al. 2015). Overall, it remains unclear whether motion priors affect perception and pursuit similarly or differently. We introduced different probabilities of motion direction in the current study, leading to an implicit expectation bias for future motion direction based on previous trial history. This manipulation allows us to investigate the effect of a general motion prior on perception and pursuit. In the following, we refer to effects of expectation as the behavior triggered by manipulations of this statistical bias.

## Methods

All three experiments were similar in terms of procedure and analyses. Experiment 1 was the main experiment, and the purpose was to compare the effect of expectation on motion direction discrimination and anticipatory pursuit. Control experiments 2 and 3 investigated alternative explanations for findings obtained in Experiment 1, testing interactions between anticipatory pursuit and perception, and effects of stimulus features, respectively. General methods are described for experiment 1. Deviations in stimuli and procedures for control experiments are briefly described in Results.

### Observers

We recruited 10 observers (age *M* = 26.20, *std* = 5.41 years; six females) with normal or corrected-to-normal visual acuity (at least 20/20 as assessed using an Early Treatment Diabetic Retinopathy Study chart) and no history of ophthalmologic, neurologic, or psychiatric disease. All observers participated in experiment 1, eight of these observers (age *M* = 27.38, *std* = 5.40 years; four females) also participated in experiment 2, and nine (age *M* = 26.56, *std* = 5.61 years; six females) also participated in experiment 3. The sample size is comparable to previous studies (*n* = 9 in Maus et al. 2015; *n* = 8 in Santos et al. 2012; *n* = 6 in Santos and Kowler 2017). The University of British Columbia Behavioral Research Ethics Board approved all experimental procedures, and all observers participated after giving written informed consent. Observers received $35 Canadian Dollars remuneration for participation per experiment.

### Visual stimuli and setup

Stimuli were random dot kinematograms (RDKs) presented in a static aperture of 20° diameter centered in the middle of the screen. Each RDK consisted of about 470 (density 1.5 dot/deg^2^) uniformly distributed white dots (98 cd/m^2^) on a grey background (22 cd/m^2^). Each dot (diameter 0.14°) moved at a constant speed of 10°/s. The dots were labeled as signal or noise dots. Labels were updated and randomly reassigned every four frames (about 47 milliseconds, ms). Signal dots always moved in the global motion direction of the RDK (left or right), while each noise dot moved in a random direction other than the signal direction with unlimited lifetime. When a dot moved out of the aperture, it reentered from the opposite side of the aperture. The coherence of the RDK was defined as the proportion of signal dots (0-100%).

Observers were seated in a dimly-lit room and viewed all stimuli on a gamma-corrected 39 cm × 29 cm CRT monitor (ViewSonic G255f; resolution 1280 × 1024 pixel; refresh rate 85 Hz). The viewing distance was 55 cm. Each observer’s head was stabilized using a chin-and-forehead-rest. Stimuli and procedure were programmed in MATLAB R2018b (The MathWorks Inc., Natick, MA) and Psychtoolbox Version 3.0.12 (Brainard 1997; Kleiner et al. 2007; Pelli 1997).

### Procedure and design

**Figure 1** shows the trial timeline in experiment 1. Observers were asked to fixate the center of the screen when the red fixation point was on for 600-900 ms. Fixation was monitored online: if eye position was further than 2° from the center of the fixation point, the fixation point turned white and the countdown of fixation duration was paused until the observer regained accurate fixation. A blank screen (gap; 300 ms) was shown to help induce anticipatory pursuit (Krauzlis and Miles 1996). Observers were then asked to smoothly follow the global motion of the RDK (700 ms) with their eyes. A dynamic white-noise mask with luminance noise randomly assigned pixel by pixel (luminance range within 7 cd/m^2^ to 46 cd/m^2^) was shown after RDK offset for 600 ms to reduce potential motion aftereffects. At the end of each trial, they were asked to report whether it moved left or right using the “left” or “right” arrow keys on the computer keyboard.

**Figure 1.**
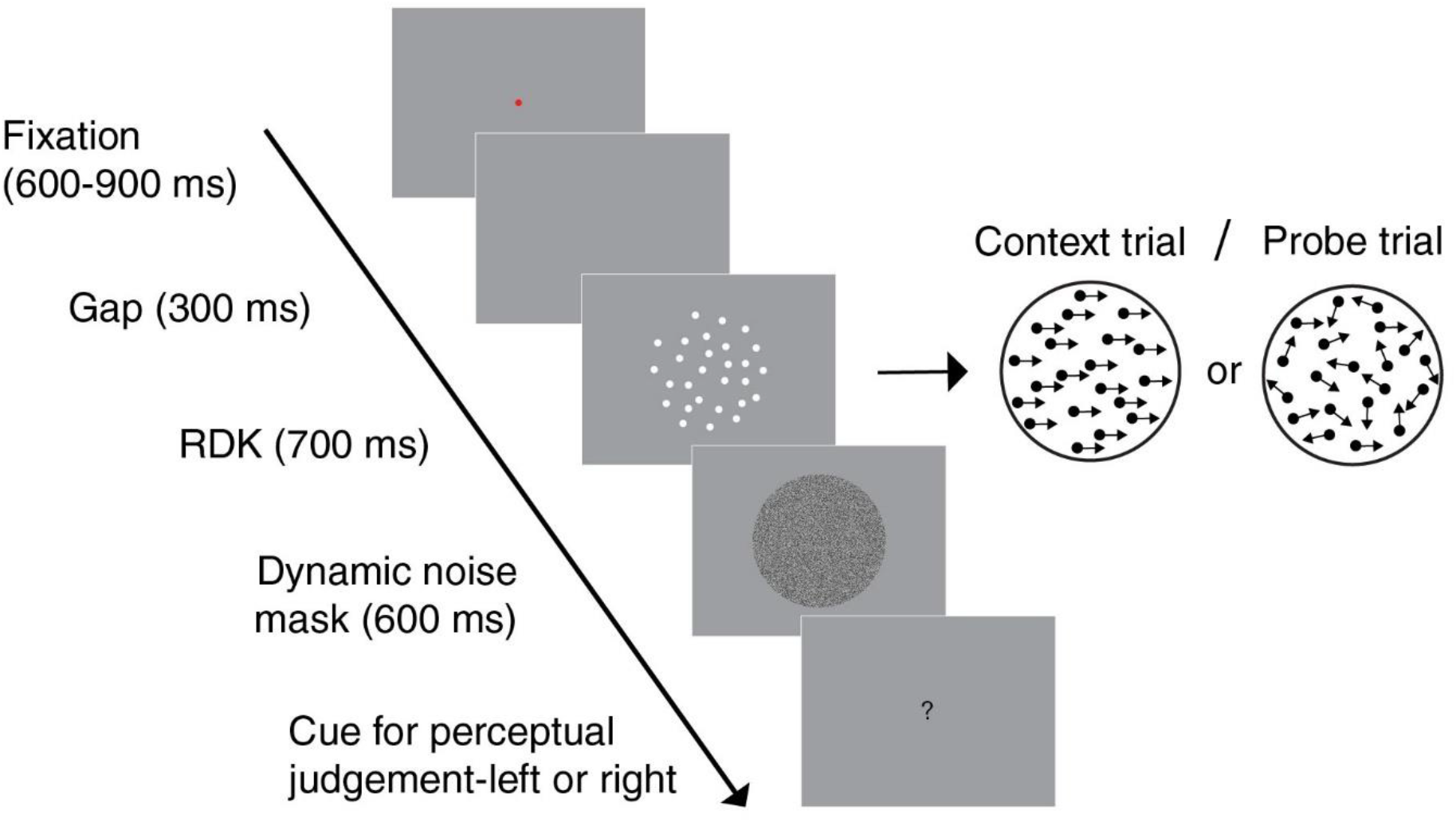
Trial timeline in experiment 1. A fixation point was shown for 600-900 ms, followed by a blank screen for 300 ms, and the RDK for 700 ms. A dynamic white-noise mask was presented after the RDK for 600 ms. Two types of trials were presented: context trials with highest-coherence RDKs, and probe trials with low-coherence RDKs. The stimuli’s relative size and contrast are increased for presentation purposes.

In each block, two types of trials were shown: 500 context trials containing non-ambiguous motion direction (100% coherence) were randomly interleaved with 182 probe trials containing ambiguous motion (0, 5, 10 or 15% coherence). The purpose of the context trials was to build up an expectation of motion direction in a given block. In each of three blocks of trials, presented in random order, we introduced different probabilities of motion direction in context trials, either 50%, 70%, and 90% probability of rightward or leftward motion. Blocks with higher probability of rightward and leftward motion were presented in different sessions; half of our sample of observers (*n* = 5) saw only higher probabilities of rightward motion, the other half saw only higher probabilities of leftward motion. The experiment was split up that way to reduce workload for each observer, because each block of trials took 60 minutes to complete for a total of 3 hours per observer. The first 50 trials in each block were always context trials. The purpose of the probe trials was to measure the effect of expectation on perception of motion direction, which would be prominent when visual input provided little evidence. In order to fairly compare perception and oculomotor anticipation we also analyzed anticipatory pursuit in probe trials only. For all observers, probe trials consisted of equal numbers of leftward/rightward trials.

### Eye movement recording and analysis

In all three experiments, the position of the right eye was recorded using a video-based eye tracker at a sampling rate of 1000 Hz (EyeLink 1000 desk-mounted, SR Research Ltd., Kanata, ON, Canada). Eye movements were then analyzed offline using custom-made MATLAB functions. Eye position, velocity, and acceleration data were filtered with a second-order Butterworth filter (cutoff frequencies of 15 Hz for position and 30 Hz for velocity and acceleration). Saccades were detected based on an acceleration criterion: the acceleration trace was segmented by zero-crossing points, and peak acceleration within each segment was calculated. If at least two consecutive segments had absolute peak acceleration larger than 400°/s^2^, these segments were defined as saccades. An acceleration threshold was used to accurately detect saccades of small amplitude and velocity during the anticipatory pursuit phase. Saccade detection was confirmed by visual inspection of the velocity traces in each trial. Saccades were then excluded from the analysis of smooth pursuit. Following previous studies (Maus et al. 2015; Santos and Kowler 2017; Watamaniuk et al. 2017), anticipatory pursuit velocity was defined as the average horizontal eye velocity during the time window from 50 ms before to 50 ms after RDK onset. We also analyzed eye velocity gain (eye velocity relative to target velocity) during visually-guided pursuit, calculated during the time window from 300 to 600 ms after target onset. Trials with blinks during RDK presentation were manually labeled as invalid and excluded (1% across observers in experiment 1, 0.5% in experiment 2, and 0.7% in experiment 3). Leftward direction is negative by convention.

### Perceptual response analysis

We did not observe systematic differences between the effects of rightward and leftward motion probability on the magnitude of anticipatory pursuit (experiment 1: *t*(4) = 0.76, *p* = .49, Cohen’s *d* = 0.94) or perceptual bias (experiment 1: *t*(4) = 1.07, *p* = .34, Cohen’s *d* = 0.83) and therefore merged data for different motion directions, presenting data as if higher probabilities of rightward motion were presented. Under each probability condition for each observer, we fitted a psychometric curve using the logistic function as shown below:

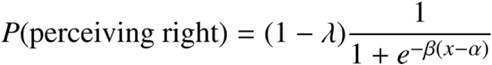

Specifically, *x* is the signed motion coherence of RDK (negative for leftward motion), α is the point of subjective equality (PSE) where observers reported both motion directions equally often (50% of the time), *β* is the slope representing the sensitivity of perception, and *λ* is the lapse rate (restricted to below 0.1 when fitting). In this study, a shift of the PSE across probability conditions would indicate a perceptual bias. A shift to the left indicates a perceptual attraction bias (i.e. with direction judgments being attracted toward the direction expectation of rightwards), and a shift to the right indicates a repulsion bias. A change in slope across probability conditions would indicate a change in sensitivity of perceptual judgments, where a steeper slope corresponds to higher sensitivity. Curve fitting was performed using the Palamedes toolbox version 1.9.0 in MATLAB (Prins and Kingdom 2018).

### Hypotheses and statistical analysis

In experiment 1, we aimed to test the following hypotheses. First, anticipatory pursuit is affected by direction expectation: the velocity of anticipatory pursuit scales positively with the target’s direction probability (attraction bias); a higher probability of rightward motion will lead to a higher velocity in anticipatory pursuit (**Fig. 2a**). Second, direction perception is affected by direction expectation: observers preferentially perceive the expected motion direction (attraction bias; **Fig. 2b**). Alternatively, perception could be biased away from the expected direction (repulsion bias; **Fig. 2b**), as has sometimes been reported in the literature. We further examined if expectation affected slope to investigate whether different prior probabilities might result in differences in sensitivity. To examine the expected effects of probability on anticipatory pursuit velocity, the magnitude of perceptual bias (shift of the PSE), and the sensitivity of perception (slope), we used one-way repeated-measures analyses of variance (ANOVA) with *probability* as factor. In addition, to examine whether anticipatory pursuit velocity and the strength of any potential perceptual bias were correlated across conditions and observers, we fitted a linear mixed-effects model of PSE with *probability* and *anticipatory pursuit velocity* as fixed effects, and individual intercept as the random effect (formula: PSE ~ anticipatory pursuit velocity + probability + (1 | observer)). Finally, we also examined the potential link between visually-guided pursuit and perception, and effects of probability on the velocity gain of visually-guided pursuit. Experiments 2 and 3 investigated alternative explanations of findings obtained in experiment 1. Their logic and underlying hypotheses are described in Results.

**Figure 2.**
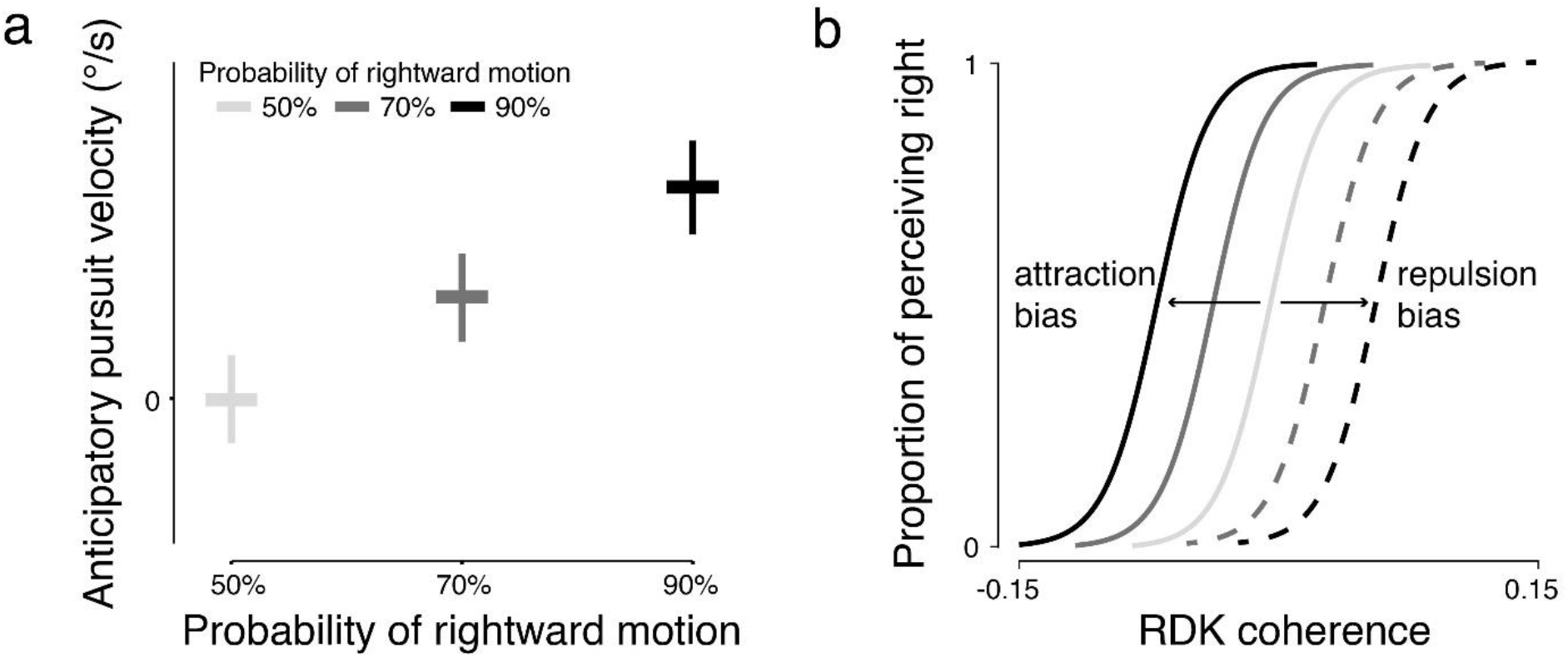
Hypotheses for (a) attraction bias in anticipatory pursuit velocity and (b) attraction or repulsion bias in perception. (a) Anticipatory pursuit velocity increases with increasing probability of rightward motion in each block, reflecting an attraction bias. (b) The perceptual bias is reflected by a shift of the PSE at higher probabilities (70% and 90%) when compared to the 50% probability condition; a leftward shift (solid lines) indicates an attraction bias, a rightward shift (dashed lines) indicates repulsion bias. Negative value of RDK coherence indicates that the global motion direction is left.

Across experiments, we report generalized eta-squared 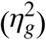 as the effect size in one-way ANOVAs, and partial eta-squared 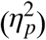 in two-way ANOVAs. For all experiments, we also report mean and 95% confidence interval (CI) of anticipatory pursuit velocity, and PSE from 1000 bootstrap simulations to supplement statistical hypothesis testing and provide quantitative estimates of the variability of sample estimates. The statistical tests were conducted in R Version 3.6.0 (package “lme4”, Bates et al. 2015; package “ez”, Lawrence 2016; R Core Team 2019) and MATLAB R2020a.

## Results

### Experiment 1

#### Evidence for attraction bias in anticipatory pursuit

Across observers and trials, we found that the velocity of the anticipatory pursuit response scaled positively with the probability of a given motion direction. **Figure 3a** shows an example of individual eye velocity traces and **Figure 3b** shows group-averaged eye velocity traces in probe trials. indicating that anticipatory pursuit velocity increases with increasing probability of rightward motion. These observations were confirmed by a significant main effect of *probability* on anticipatory pursuit velocity (**Fig. 3c**), *F*(2,18) = 28.19, *p* = 2.84×10^−6^, 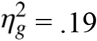. As a complementary method to these statistics, the bootstrapped mean and 95% CI of anticipatory pursuit velocity confirmed our findings (50%: 0.24 ± 0.01°/s; 70%: 0.70 ± 0.01°/s; 90%: 1.26 ± 0.01°/s).

**Figure 3.**
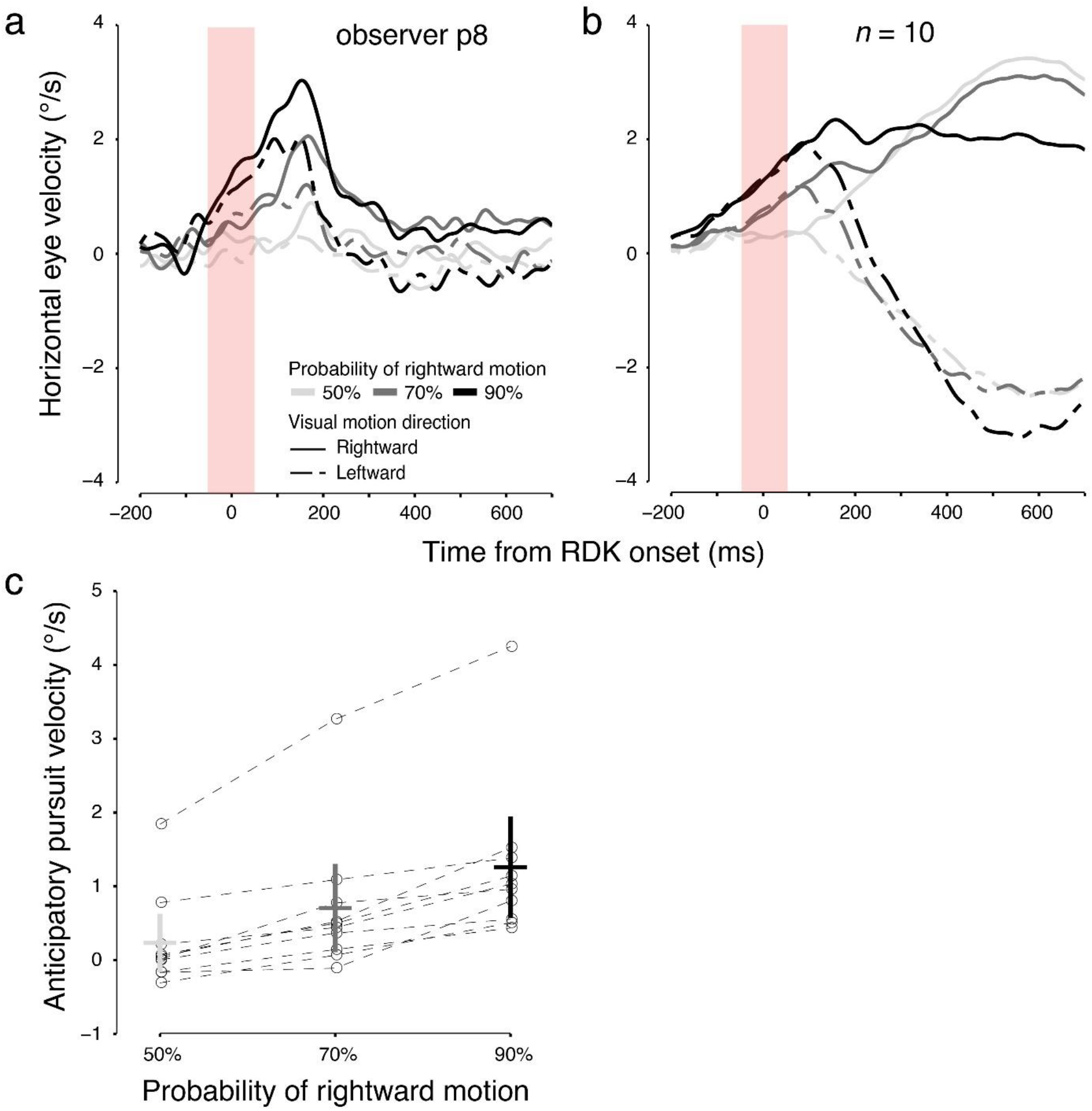
(a) Example trial-average horizontal eye velocity traces in probe trials (leftward or rightward motion direction as indicated by line type) for different probability conditions (indicated by line color) for one representative observer in experiment 1. The red shaded area indicates the analysis window for anticipatory pursuit. This particular observer had little visually-guided pursuit in probe trials. (b) Group-average horizontal eye velocity traces in probe trials for different probability conditions in experiment 1 for *n* = 10. Line types in panels and b denote motion direction and probability of rightward motion. (c) Horizontal anticipatory pursuit velocity in experiment 1 averaged across the time interval indicated as shaded area in panel a. Horizontal bars indicate the mean anticipatory pursuit velocity across observers, and vertical bars indicate the 95% CI. The circles indicate the mean anticipatory pursuit velocity of individual observers, connected by dashed lines across probability conditions. Results were the same even if excluding the one outlier who had relatively high anticipatory pursuit velocity.

#### Evidence for repulsion bias in direction perception

Perceptual results are incongruent with what we observed for anticipatory pursuit. We found a systematic rightward shift in the PSE at the individual observer level (**Fig. 4a**) as well as across observers (**Fig. 4b**), indicating a perceptual bias away from the high-probability motion direction. When rightward trials had a higher probability in context trials, observers tended to perceive leftward direction more often in probe trials. These observations were confirmed by a significant main effect of *probability* on the PSE (**Fig. 4b**), *F*(2,18) = 20.36, *p* = 2.39×10^−5^, 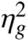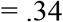. The bootstrapped mean and 95% CI of the PSE confirmed these statistical results (50%: −0.02 ± .002; 70%: 0.0003 ± 0.002; 90%: 0.04 ± 0.0002). We did not observe any significant effects of *probability* on slope (*F*(2,18) = 0.78, *p* = .48, 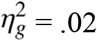), indicating that sensitivity did not change across probability conditions. The bootstrapped mean and 95% CI of slope were 31.31 ± 5.26 for 50%, 31.72 ± 2.14 for 70%, and 29.87 ± 3.93 for 90%.

**Figure 4.**
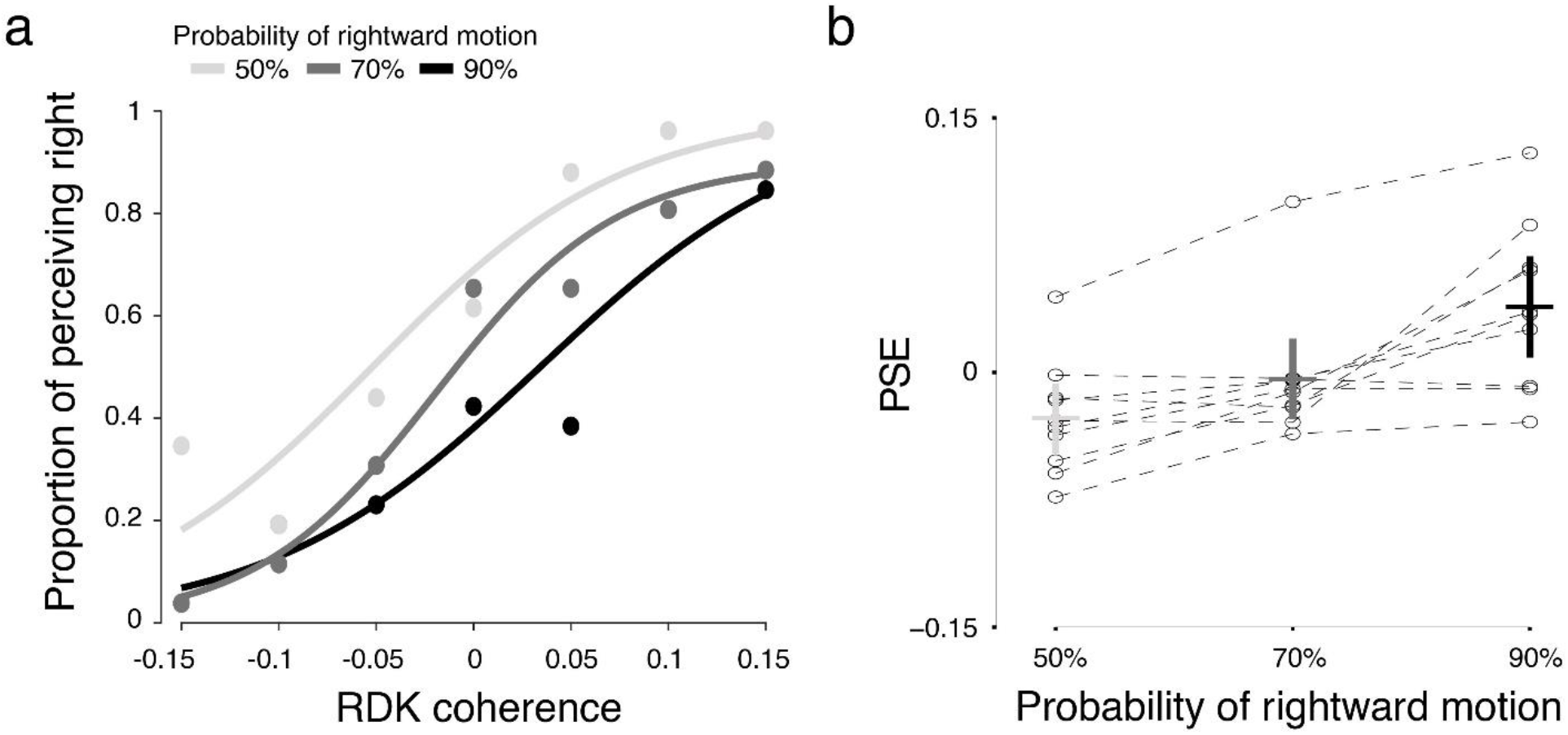
(a) Example psychometric curves from the same individual observer shown in Fig. 3a. Negative coherence represents leftward direction. (b) PSE in experiment 1 (*n* = 10).

We did not find any dependencies between PSE and anticipatory pursuit velocity in addition to the fact that they both changed with probability. The fixed effect of anticipatory pursuit velocity in the linear mixed-effects model of PSE (PSE ~ anticipatory pursuit velocity + probability + (1 | observer)) was not significant (estimate ± *std* = −0.002 ± 0.01, *t*(20.31) = −0.16, *p* = .87), and only the fixed effect of probability was significant (estimate ± *std* = 0.002 ± 0.0004, *t*(30.00) = 4.60, *p* = 7.12×10^−5^).

Taken together, our results point at a differential effect of motion direction probability on anticipatory pursuit, reflecting an attraction bias, and direction perception, reflecting a repulsion bias. To explore this further, we next examined the potential link between visually-guided pursuit and perception, and analyzed the effect of probability on visually-guided pursuit velocity gain.

#### Visually-guided pursuit is aligned with direction perception

Whereas anticipatory pursuit is mostly driven by expectation, visually-guided pursuit is tuned to the visual properties of the target and is known to strongly covary with motion perception (Spering and Montagnini 2011). Previous research has demonstrated that smooth pursuit can be elicited by perceived illusory motion rather than by physical motion (Madelain and Krauzlis 2003; Montagnini et al. 2006). Therefore, visually-guided pursuit could follow different result patterns from anticipatory pursuit and be more aligned with perception.

Here we investigated whether visually-guided pursuit was more in line with an attraction bias (as in anticipatory pursuit) or followed a repulsion bias (as in perception). We compared eye velocity gain in conditions in which the perceptual judgment corresponded to the physical motion direction (congruent) with conditions where perceptual judgments went in the opposite direction to the physical motion (incongruent). **Figure 5** shows average velocity traces in probe trials across all observers for congruent (**Fig. 5a**) versus incongruent trials (**Fig. 5b**). Note that this categorization of congruency is agnostic on whether perception followed the expected motion direction or the opposite one and merely reflects how closely perception matched physical target motion. Whereas late visually-guided pursuit followed the visual motion direction in congruent trials (shaded areas in **Fig. 5a**), pursuit followed visual motion direction less in incongruent trials (**Fig. 5b**) with a tendency to be directed into the opposite (perceived) direction, resulting in smaller or negative gains. This is confirmed by a significant main effect of *congruency* on velocity gain (**Fig. 5c**; *F*(1,9) = 20.65, *p* = .001, 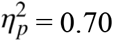). The main effect of *probability* (*F*(2,18) = 0.049, *p* = .953, 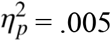) or the *congruency* × *probability* interaction (*F*(2,18) = 0.051, *p* = .950, 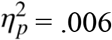) were not significant. This difference in visually-guided pursuit between congruent and incongruent conditions persisted across different levels of motion coherence (not shown).

**Figure 5.**
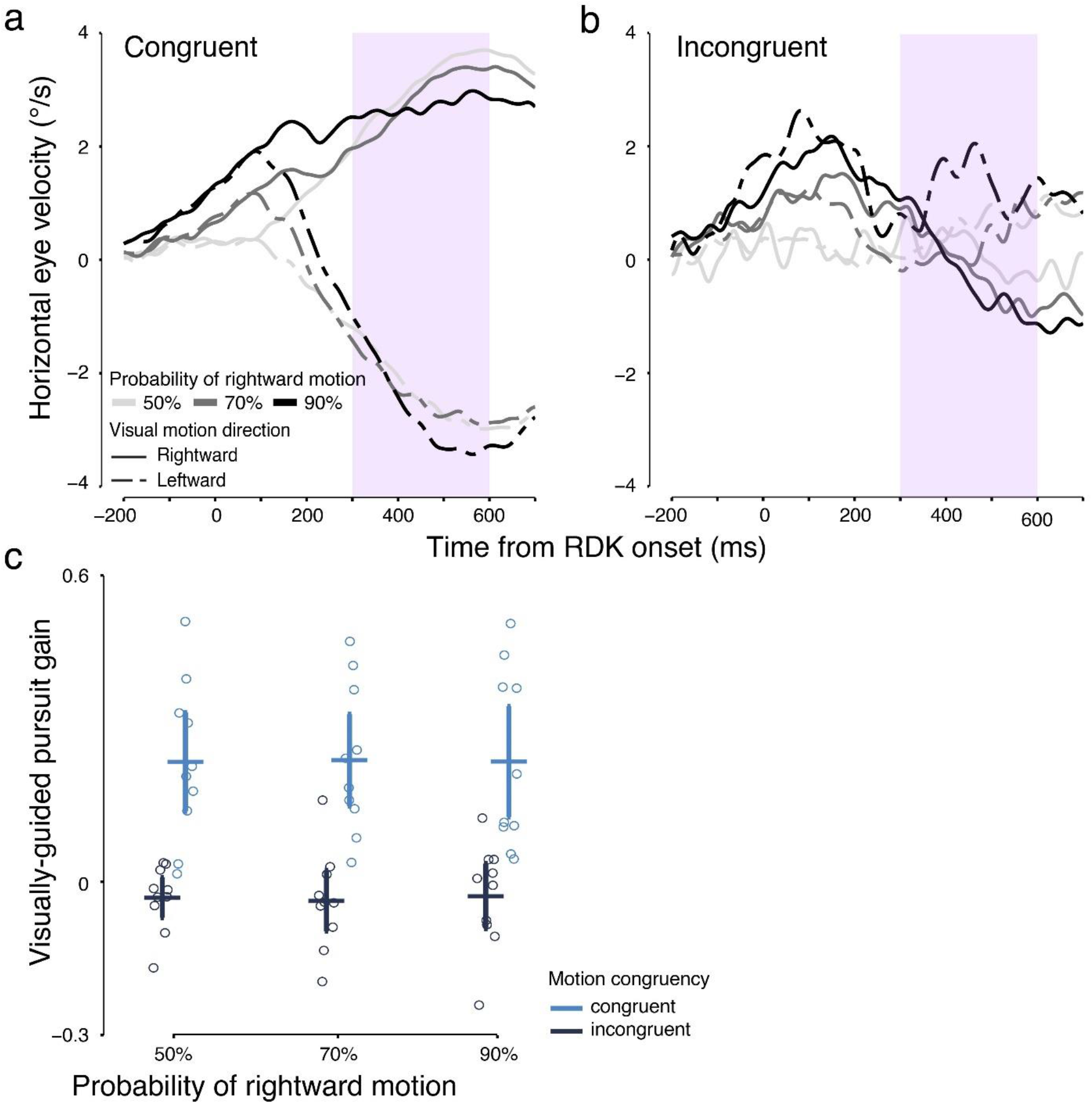
(a) Group-averaged (*n* = 10) horizontal eye velocity traces in probe trials in which visual motion directions were congruent (3967 trials total) with perceived directions across different probability conditions in experiment 1. The purple shaded area indicates the analysis window for late-phase pursuit; an early cutoff was applied to reduce the effect of anticipatory deceleration before the end of each trial (at 700 ms). (b) Group-averaged (*n* = 10) horizontal eye velocity traces in probe trials in which visual motion directions were incongruent (649 trials total) with perceived directions across different probability conditions in experiment 1. (c) Late-phase visually-guided pursuit gain in probe trials grouped by motion congruency across different probability conditions in experiment 1 (*n* = 10). Higher gain indicates that the eyes follow the visual motion better, and negative gain indicates that the eyes are moving in the opposite direction to the visual motion direction. Horizontal bars indicate the mean visually-guided pursuit gain across observers, and vertical bars indicate the 95% CI. The circles indicate the mean visually-guided pursuit gain of individual observers.

Further, we examined if expectation had an effect on visually-guided pursuit, by analyzing the effect of *probability* on visually-guided pursuit gain. Late-phase visually-guided pursuit in rightward probe trials seemed to have lower velocity in blocks with higher probability of rightward motion (see **Fig. 3b**, can also be seen in **Fig. 5a** with majority of probe trials). Since direction expectation might affect visually-guided smooth pursuit differently for rightward and leftward motion trials, we included *visual motion direction* as a second factor in the two-way ANOVA on visually-guided pursuit gain. If the observed decrease in eye velocity with increased probability across blocks was true, a significant main effect of *probability* and possibly a significant interaction effect of *probability* × *visual motion direction* could be observed. However, the interaction effect (*F*(2,18) = 1.54, *p* = .24, 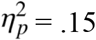; **Fig. 6**), the main effect of *probability* (*F*(2,18) = 0.58, *p* = .57, 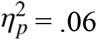), or the main effect of *visual motion direction* (*F*(1,9) = 1.87, *p* = .20, 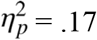) were non-significant. This could be due to the large individual variability—some observers had little visually-guided pursuit in probe trials across probability conditions (gain close to zero), likely due to the low RDK coherence.

**Figure 6.**
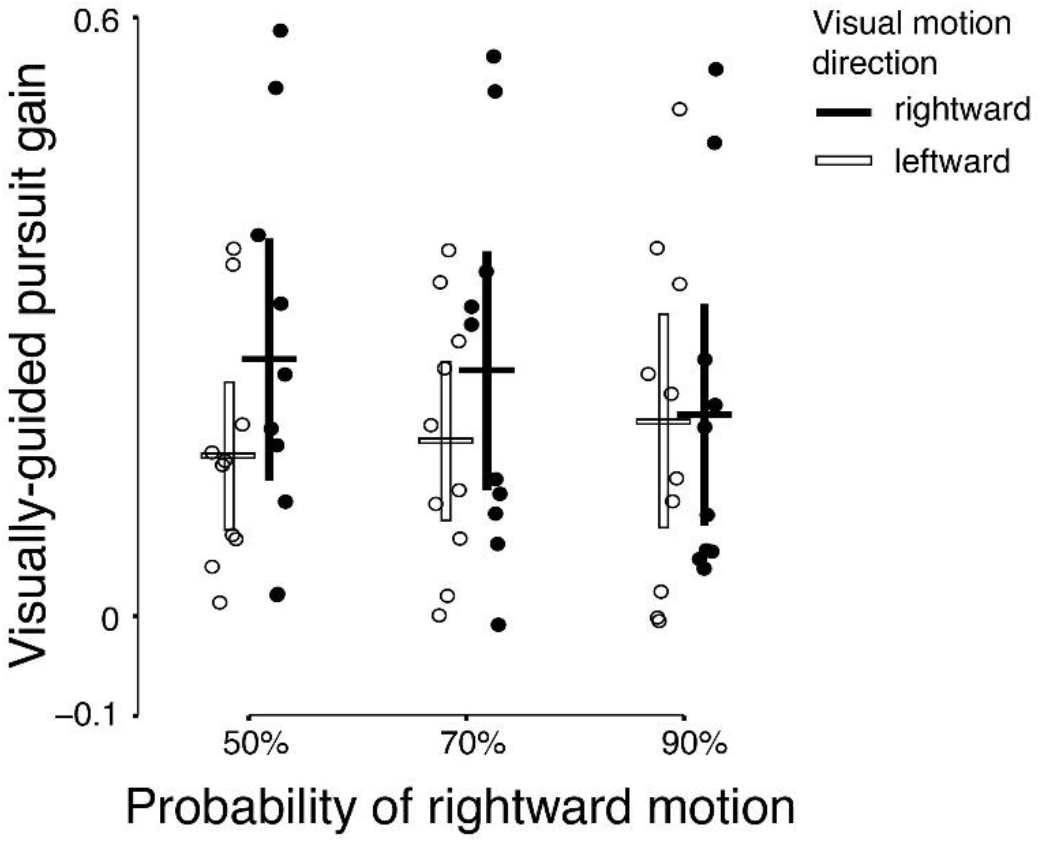
Late-phase visually-guided pursuit gain in probe trials grouped by visual motion direction across different probability conditions in experiment 1 (*n* = 10).

Notwithstanding a lack of evidence for expectation effects on visually-guided pursuit, this part of the pursuit response was more aligned with perception than anticipatory pursuit. This finding indicates that late-phase pursuit is driven by signals that are more coherent with those signals driving perceptual judgments than those driving anticipatory pursuit.

### Experiment 2

To further explore the mechanisms underlying the dissociation between expectation effects on anticipatory pursuit and perception, we conducted two control experiments. One potential problem with our paradigm might be that anticipatory pursuit during the earliest phase of the presentation of the low-coherence RDK elicits retinal image motion in the opposite direction than the expected one. This motion signal could have informed the perceptual choice, explaining the repulsion bias. In experiment 2, we therefore tested whether the observed perceptual bias was affected by this negative retinal motion signal by manipulating anticipatory pursuit magnitude. To reduce anticipatory pursuit, we showed the fixation point until RDK onset, omitting the 300-ms gap introduced in Experiment 1, and instructed observers to maintain fixation until the stimulus started moving. Each observer completed two blocks (50% and 90% probability of rightward motion). All other procedures were the same as in experiment 1.

To confirm that anticipatory pursuit was reduced in Experiment 2 as compared to Experiment 1, we performed a two-way repeated measures ANOVA with *experiment* and *probability* as factors. An *experiment* × *probability* interaction effect on anticipatory pursuit velocity would indicate a change in anticipatory pursuit magnitude from one experiment to the other. If anticipatory pursuit induced the perceptual bias, reduced anticipatory pursuit magnitude in Experiment 2 should result in a smaller perceptual bias. This interpretation would be supported by a significant *experiment* × *probability* interaction effect on PSE.

#### Anticipatory pursuit was significantly reduced with prolonged fixation

The experimental manipulation of prolonging fixation in Experiment 2 yielded the expected reduction in anticipatory pursuit velocity from 1.26 ± 1.11°/s (*M* ± *std*) in Experiment 1 to 0.57 ± 0.30 °/s in Experiment 2 at the highest probability of rightward motion (**Fig. 7a, b)**. This observation was confirmed by a significant *experiment* × *probability* interaction (**Fig. 7c;** *F*(1,7) = 7.20, *p* = .03, 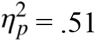). Despite lower overall velocity, higher probability of rightward motion continued to induce higher anticipatory pursuit velocity, reflected in a main effect of *probability* (*F*(1,7) = 37.81, *p* = .0005, 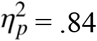). Congruently, the bootstrapped mean and 95% CI of anticipatory pursuit velocity in experiment 2 were .05 ± .01°/s for 50% and .59 ± .01°/s for 90%. The main effect of *experiment* was not significant (*F*(1,7) = 2.06, *p* = .19, 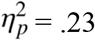).

**Figure 7.**
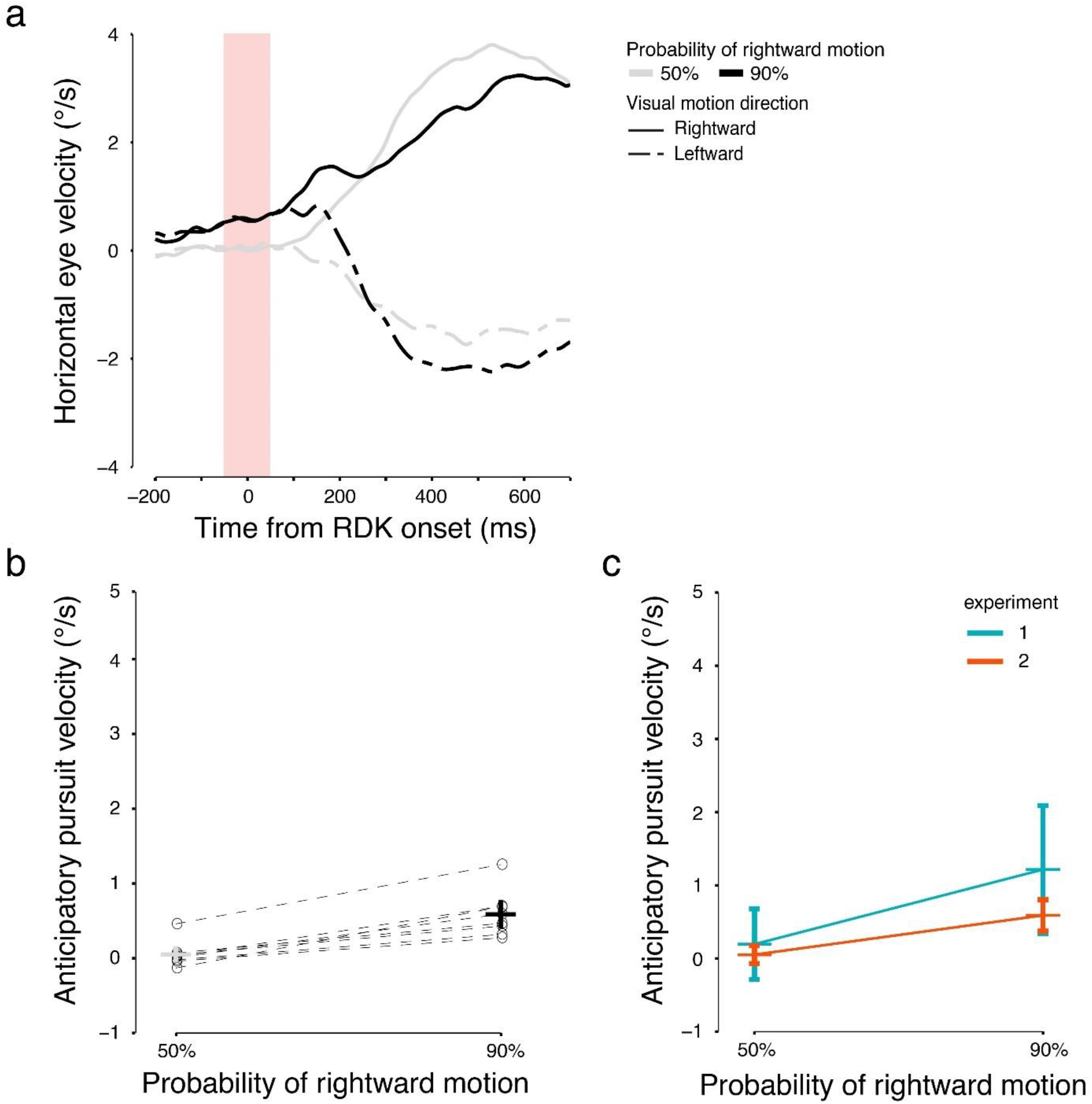
(a) The group-averaged (*n* = 8) horizontal eye velocity traces in probe trials across different probability conditions in experiment 2. (b) Horizontal anticipatory pursuit velocity in experiment 2 (*n* = 8). (c) The comparison of anticipatory pursuit velocity between experiment 1 and 2. The horizontal bars show the mean across observers (*n* = 8 for both experiments), and the error bars show the 95% CI.

#### Persistent perceptual bias despite reduced anticipatory pursuit velocity

Despite the successful reduction in anticipatory pursuit velocity, we observed the same repulsion bias (rightward shift of the PSE; **Fig. 8a**) in perceptual judgments in Experiment 2. This observation was confirmed by a significant main effect of *probability*, *F*(1,7) = 22.91, *p* = .002, 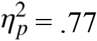, and a lack of *experiment* × *probability* interaction (**Fig. 8b**; *F*(1,7) = 0.001, *p* = .97, 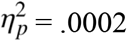). The bootstrapped mean and 95% CI of the PSE were −.04 ± .003 for 50% and .02 ± .002 for 90%. The main effect of *experiment* was not significant (*F*(1,7) = 4.54, *p* = .07, 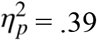). These results indicate that the perceptual bias remained stable across experiments, and that negative retinal image motion induced by anticipatory pursuit is unlikely to cause the repulsion bias.

**Figure 8.**
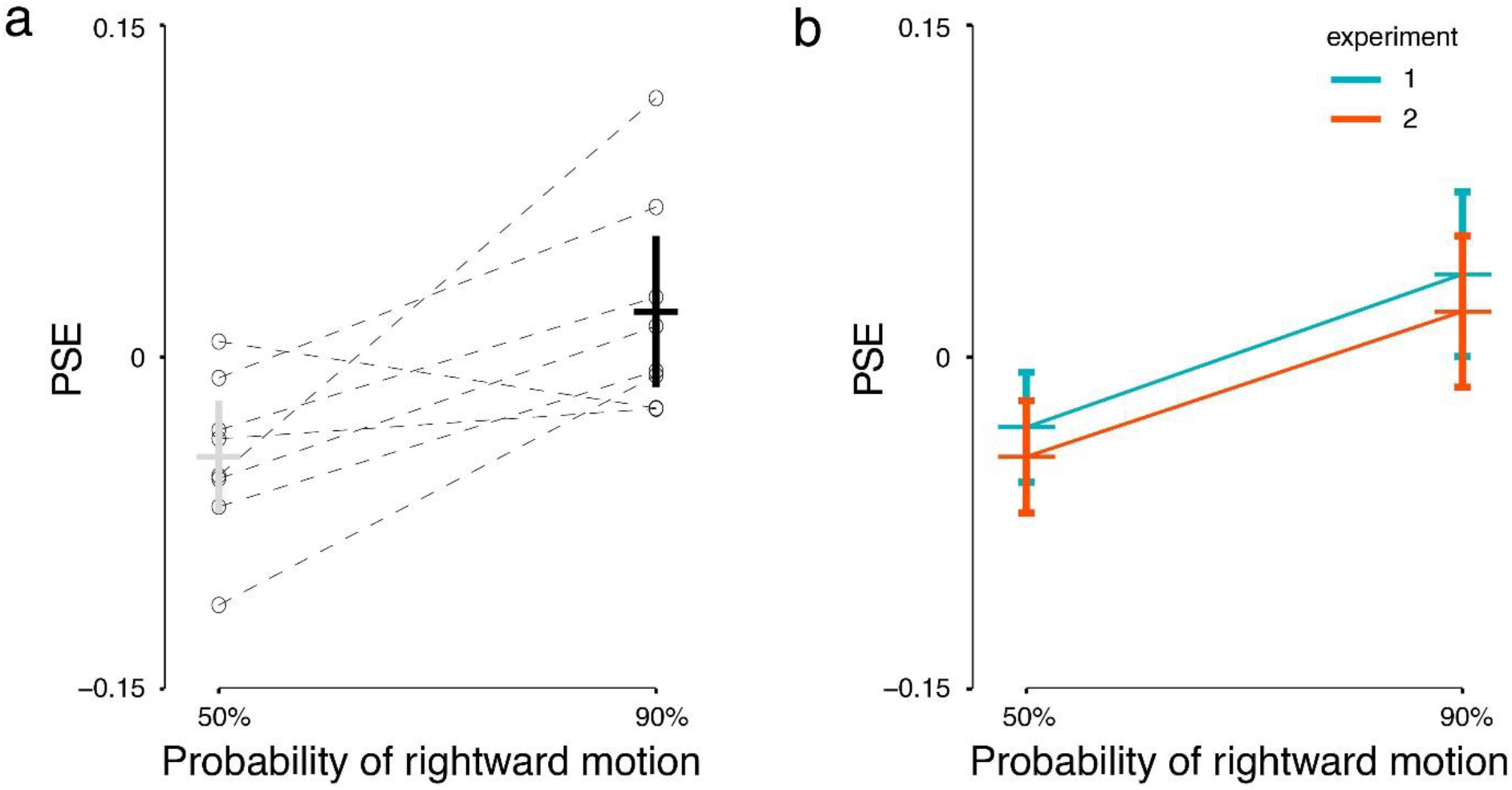
(a) PSE in probe trials across different probability conditions in experiment 2 (*n* = 8). (b) The comparison of PSE between experiment 1 and 2 (*n* = 8 in each experiment).

### Experiment 3

In all experiments presented so far, we used a noise mask following RDK presentation (**Fig. 1**) to reduce potential motion aftereffects. Such aftereffects have been observed in perception (Mather et al. 2008) and pursuit (Braun et al. 2006). An alternative explanation for the perceptual repulsion effect observed in experiments 1 and 2 could be that prolonged exposure to a high-coherence moving stimulus in context trials produces a perceptual aftereffect (a form of low-level sensory adaptation) despite the mask. One way to reduce potential effects of motion aftereffects or other similar forms of sensory adaption is to lower the motion signal strength of the adaptor, for example, by reducing its luminance contrast (Keck et al. 1976). It is well known that the response of neurons in motion-sensitive middle temporal cortex (area MT) is modulated by motion coherence (Händel et al. 2007). In Experiment 3, we therefore reduced the coherence of the RDK in context trials to investigate whether such a manipulation would weaken the perceptual repulsion bias. We reduced motion coherence of RDKs in context trials to 25% on average (coherence levels of 20% and 30% randomly assigned to half of the context trials in each block). This coherence level is considered to be above perceptual thresholds for direction discrimination in adults (Meier and Giaschi 2014) and yielded judgements of >99% correct accuracy in context trials in our experiment. We therefore expected that the perceived probability of context trials (50% and 90%) remained the same as in previous experiments. All other procedures were the same as in experiment 1.

First, we assessed whether coherence impacted visually-guided pursuit in context trials to confirm that the coherence manipulation successfully reduced the motion signal. We conducted a two-way repeated-measures ANOVA on pursuit gain with *experiment* and *probability* as factors. A significant main effect of *experiment* would imply a reduction in motion signal due to the reduced coherence. To examine if motion coherence has an effect on anticipatory pursuit, we conducted a two-way repeated-measures ANOVA on anticipatory pursuit velocity with *experiment* and *probability* as factors. A significant interaction would indicate that anticipatory pursuit was modulated by motion signal strength in context trials. Second, to examine whether RDK coherence in context trials affects perception, we conducted a two-way repeated-measures ANOVA on PSE with *experiment* and *probability* as factors. If RDK coherence in context trials affected the repulsion bias, we should find a significant interaction.

#### Low-coherence context trials elicit weaker visually-guided and anticipatory smooth pursuit

The experimental manipulation of motion coherence yielded the expected reduction in visually-guided pursuit gain in context trials (exp. 1: *M* = 0.89 ± 0.11 across observers and probability conditions, exp. 3: *M* = 0.49 ± 0.16). This observation was confirmed by a significant main effect of *experiment* on pursuit gain (*F*(1,8) = 192.62, *p* = 7.03 × 10^−7^, 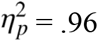). No significant main effect of *probability* (*F*(1,8) = 1.66, *p* = .23, 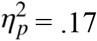) or interaction (*F*(1,8) = 1.68, *p* = .23, 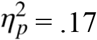) were found. This confirmed that a reduction in motion coherence elicited a weaker motion signal and therefore lower pursuit gain in context trials.

Similarly, the experimental manipulation of motion coherence in context trials reduced anticipatory pursuit velocity at the highest probability of rightward motion in probe trials (exp. 1: *M* = 1.26 ± 1.11°/s; exp. 3: *M* = 0.62 ± 0.62°/s, **Fig. 9a, b**). This observation was confirmed by a significant *experiment* × *probability* interaction effect (*F*(1,8)=32.39, *p*=.0005, 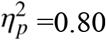), indicating that the effect of *probability* was smaller in experiment 3 than in experiment 1 (**Fig. 3c**). The main effect of *probability* was also significant (*F*(1,8) = 23.29, *p* = .001, 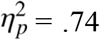), and the main effect of *experiment* was not significant (*F*(1,8) = 5.10, *p* = .05, 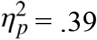). The bootstrapped mean and 95% CI of anticipatory pursuit velocity in experiment 3 were 0.17 ± 0.01°/s for 50% and 0.61 ± 0.01 °/s for 90%.

**Figure 9.**
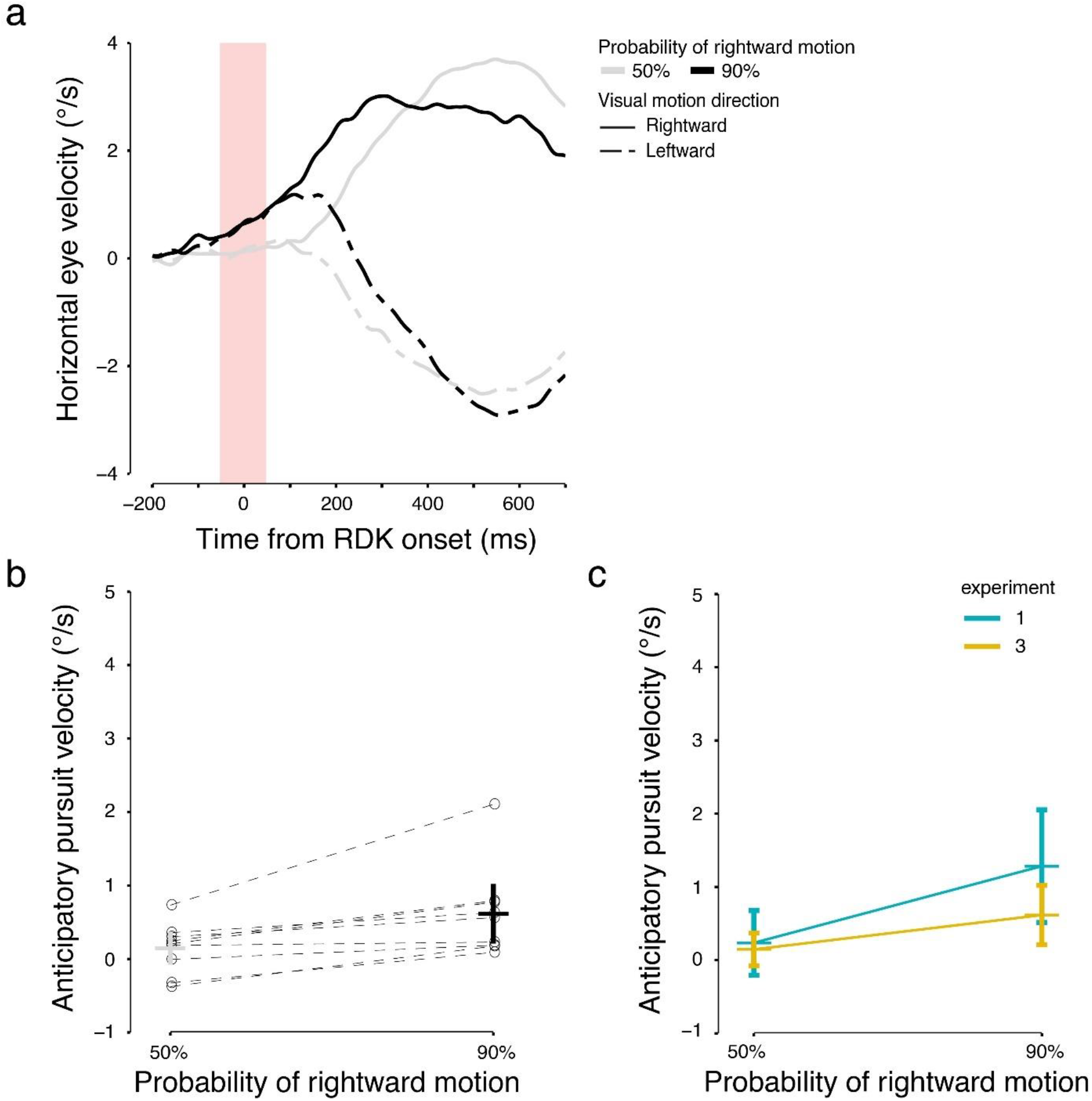
(a) The averaged (*n* = 9) horizontal eye velocity traces in probe trials across different probability conditions in experiment 3. (b) Horizontal anticipatory pursuit velocity in experiment 3 (*n* = 9). (c) The comparison of anticipatory pursuit velocity between experiment 1 and 3 (*n* = 9 in each experiment).

#### Persistent perceptual bias despite reduced motion coherence

Even though we found a reduction in visually-guided pursuit velocity, confirming that a lower-coherence stimulus shown in context trials elicited weaker pursuit, and thus was less likely to cause sensory adaptation, the perceptual bias remained stable (**Fig. 10**). This observation was confirmed by a lack of *experiment* × *probability* interaction on the PSE (*F*(1,8) = 0.53, *p* = .49, 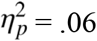), indicating a similar magnitude of perceptual bias in both experiments (**Fig. 10**). The main effect of *probability* was significant (*F*(1,8) = 44.97, *p* = .0002, 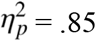), and the main effect of *experiment* was not significant (*F*(1,8) = 0.13, *p* = .73, 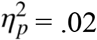). Congruently, the bootstrapped mean and 95% CI of the PSE were −.02 ± .002 for 50% and .05 ± .002 for 90%.

**Figure 10.**
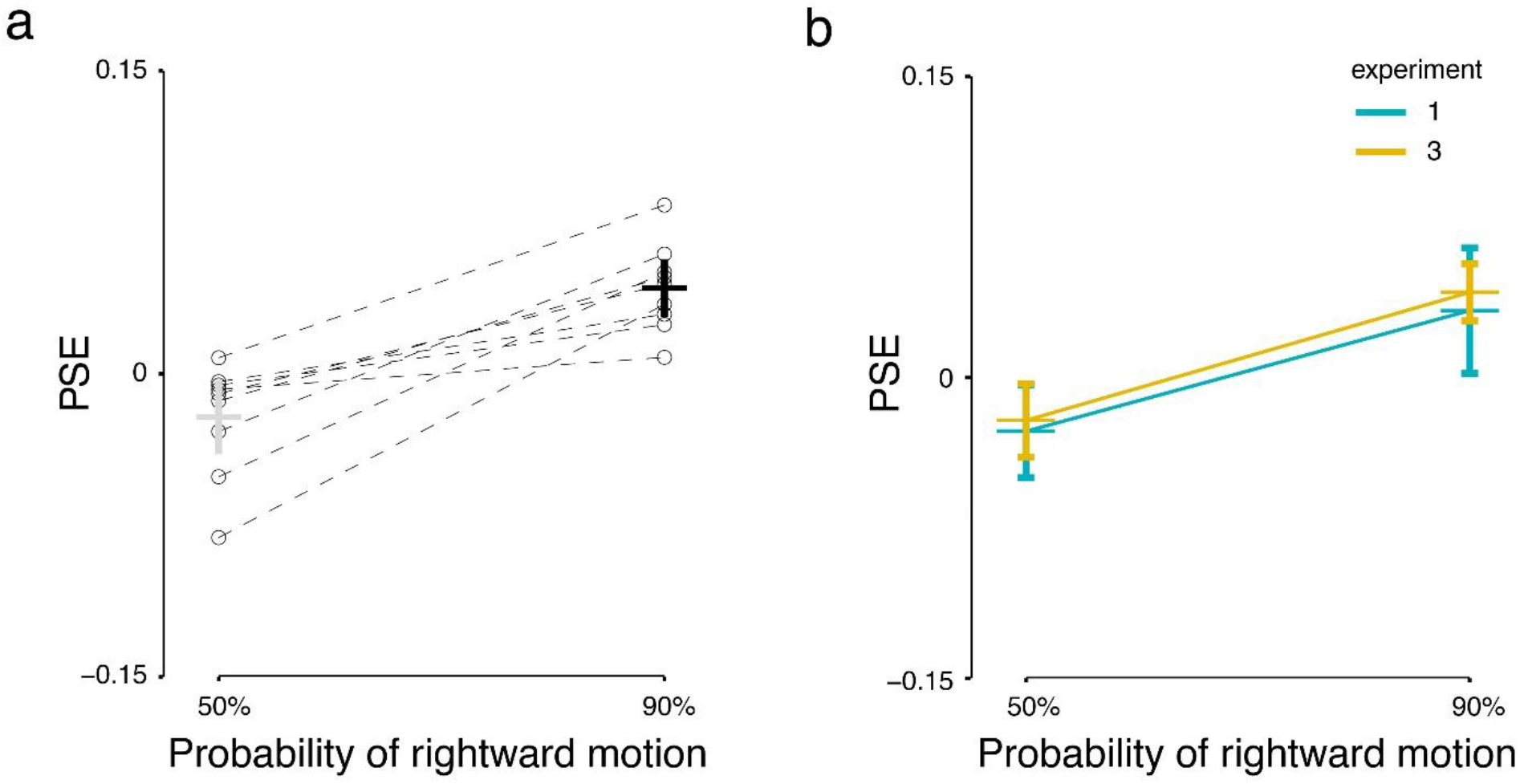
(a) PSE in probe trials across different probability conditions in experiment 3 (*n* = 9). (b) Comparison of PSE between experiment 1 and 3 (*n* = 9 in each experiment).

In summary, results from experiment 3 suggest that the perceptual bias was not purely an aftereffect induced by repeated exposure to strong motion signals, because reduced motion coherence in the context trial history did not modulate the perceptual repulsion bias.

## Discussion

By introducing a prior based on different probabilities of rightward motion, we tested how expectation affected anticipatory pursuit and direction perception. Anticipatory pursuit was directed toward the highest probability direction in a given block of trials. By contrast, the direction of perceptual judgments was repelled away from the most probable direction. This repulsion bias was not caused by anticipatory pursuit (experiment 2) and unlikely to be caused by sensory adaptation (experiment 3).

These results make a novel contribution to the literature on the comparison between perception and pursuit and highlight that motion perception does not necessarily rely on expectation in the same way as early, anticipatory pursuit responses do. These findings are generally congruent with a previous study investigating trial history effects in the velocity domain (Maus et al. 2015). In that study, the authors presented random sequences of brief, dot-motion stimuli with different speeds. Observers were asked to track the motion with their eyes and judge whether the current trial’s speed was faster or slower than the speed averaged across previous trials. Whereas anticipatory pursuit scaled with previous target speed, perceptual judgments were faster after slow stimuli and vice versa for fast stimuli, indicating a similar repulsion effect as observed for motion direction in the current study. Taken together, our study and the previous study show that opposite effects on perception and anticipatory pursuit exist regardless of which feature of the stimulus is manipulated (speed vs. direction) and over which time course both responses are compared (long-term prior vs. shorter-term trial history).

Such opposite biases and different sensitivities to manipulations of motion signal strength in perception and pursuit are generally compatible with the idea of different information processing for both responses. Perception and visually-guided pursuit eye movements might rely on different information accumulation and integration over time, due to different needs of the perceptual and the oculomotor systems (Spering and Montagnini 2011). The current paper mainly compares perceptual responses to another aspect of pursuit, the earliest, anticipatory phase that is driven by cognitive, memory-related signals rather than by visual signals. Interestingly, the present study did not find a significant effect of direction expectation on the later, visually-guided phase of smooth pursuit. However, in trials in which the perceived direction was incongruent with the physical motion direction (i.e., followed a repulsion bias), visually-guided smooth pursuit was aligned with perception. In the following paragraphs, we will discuss the characteristics of the signals driving motion perception and different phases of smooth pursuit with a main focus on anticipatory pursuit.

### Different biases in motion perception and pursuit might reflect how both responses adapt to task requirements

Studies on perceptual responses to manipulations in short-term or long-term probability have mostly revealed an attraction bias (Alais et al. 2017; Chalk et al. 2010; Kok et al. 2013), in which perception follows the recently viewed or most likely stimulus feature (direction, orientation, etc.). Our paradigm utilized short-duration displays and introduced a statistical bias of motion direction (long-term probability), similar to some of these studies (e.g., Chalk et al. 2010). However, we observed a repulsion bias. These different types of biases are interesting because they reveal that the perceptual system might respond to different task and stimulus environments in a flexible way, depending on the requirements of the task (e.g., to sensitively respond to a change or categorize information). A repulsion bias might reflect the need of the perceptual system to stay alert and to quickly respond to changes in the environment in an energy-efficient way: there is no need to be highly sensitive to a stimulus that always appears, whereas a novel stimulus would alert the system and may require priority processing. This is similar to the functional role of adaptation, yet results from experiment 3 suggested that the repulsion perceptual bias in our experiment was not caused by low-level sensory adaptation.

Similarly, different result patterns in perception and pursuit might reflect different task requirements as well. Our results resemble those obtained in other studies comparing perception and pursuit (Spering and Gegenfurtner 2007), or pursuit and hand movements (Kreyenmeier et al. 2017). These studies found that pursuit generally followed the motion average of different target and context speeds, whereas perception and manual interception of a target followed the difference of target and background. Akin to the task requirements in our study, perception and action served different functions in these studies as well. Whereas perception’s role appeared to be to segregate a target from the background, pursuit’s role was to track in the general target direction in the presence of surrounding motion signals, resulting in motion vector averaging. In the present work the tendency for pursuit to rely on integration rather than segmentation of different signals seems to be extended to the trial-history domain, and to the anticipatory phase: integration of visual signals across many trials would drive anticipatory pursuit, whereas perception could be based on the contrast between the current visual input and long-term memory of visual motion.

### Expectation effects on anticipatory vs. visually-guided pursuit

Anticipatory pursuit has been shown to be not purely habitual (Jarrett and Barnes 2001, 2002; Kowler 1989), but also sensitive to different types of cognitive cues and probabilistic context (Pasturel et al. 2020; Santos and Kowler 2017) as well as to reward (Damasse et al. 2018). In addition, we show that anticipatory pursuit velocity is modulated by motion coherence in context trials, i.e. motion signal strength in the prior history. Our results together with previous findings suggest that anticipatory pursuit is based on the integration of multiple signals, from low-level visual motion signals (weighted by the sensory strength, or saliency, e.g. RDK coherence), to higher-level cognitive cues such as expectation and reward. This holds across the short time-scale of a single trial for visually-guided early smooth pursuit (e.g., Ferrera and Lisberger 1997) and a much longer time-scale lasting several minutes for anticipatory pursuit, as in our experimental blocks. The assumption that anticipatory pursuit is based on such an integration of multiple signals is reasonable considering that the goal of anticipatory pursuit is to reach accurate tracking as soon as possible, in order to reduce the temporal delay in tracking the visual target.

Interestingly, whereas anticipatory pursuit showed an attraction bias scaling with probability, late-phase visually-guided pursuit did not follow the same result pattern. This difference might be expected given that visually-guided pursuit and perception similarly rely on current sensory signals, whereas anticipatory pursuit is driven by expectation. Moreover, it is known that visually-guided pursuit and motion perception interact, and that motion perception can modulate pursuit (Madelain and Krauzlis 2003; Montagnini et al. 2006). It is thus possible that the late-phase visually-guided pursuit observed in our experiment was driven by perception, regardless of the nature of the expectation information.

### Neural correlates of expectation effects on perceptual bias and anticipatory pursuit

The dissociation between effects of expectation on perception and anticipatory pursuit might be due to perception and pursuit depending on different cortical areas during different processing stages. For perception, modulation by expectation might have affected sensory processing of current stimuli, based on activity in early visual cortical areas. By contrast, anticipatory pursuit is not triggered by current stimuli but instead based on expectation or history, related to activity in frontal cortical areas. In the following paragraphs we will discuss the neural correlates for perception and anticipatory pursuit accordingly.

Cortical area MT and the medial superior temporal area (MST) are the major sensory areas for motion processing for both perception and visually-guided smooth pursuit (Born and Bradley 2005; Thier and Ilg 2005). It is unclear which specific cortical areas are responsible for expectation effects on perception. However, there is evidence that modulation in early sensory cortices, from primary visual cortex (V1) to MT and MST, might underlie repulsion and attraction biases in perception. For example, the repulsion bias in perceived orientation was found to be stronger when the current and previous stimuli in a given trial were presented at the same location (Fritsche et al. 2020). These findings indicate that this orientation bias was driven by effects that are spatially specific (and retinotopically congruent), which likely implies modulation of neurons in early sensory cortex, such as V1, responsive to stimuli within small receptive fields. Similarly, spatial specificity has been found in visual motion adaptation in relation to neuronal activity in area MT (Kohn and Movshon 2003). Area MT is known for its large receptive fields and might inherit spatial specificity from V1, but also shows distinct adaptation responses (Kohn and Movshon 2004) that could underlie a repulsion bias. Although our results indicate that such low-level sensory adaptation was unlikely the cause of the repulsion bias in our study, other mechanisms could have resulted in a similar modulation of MT neuronal activity, leading to a repulsion bias in perception. Congruently, for attraction biases induced by expectation, functional magnetic resonance imaging studies (e.g., Kok et al. 2013) showed that representation of visual motion direction in early sensory cortices from V1 to MT was biased toward the expected stimulus. Taken together, these studies indicate that sensory areas as early as V1, and potentially up to MT and MST, play an important role in the effect of expectation on motion perception. Whether the modulation on early sensory cortex comes from higher-level areas remains unclear.

The supplementary eye field (SEF) plays a critical role for anticipatory pursuit, as shown in studies using electrical microstimulation in SEF to elicit anticipatory pursuit (Missal and Heinen 2004). Moreover, direction-selective neurons in area SEF showed stronger activity before anticipatory pursuit in their preferred direction, indicating that SEF plays a role in the preparation of anticipatory pursuit (de Hemptinne et al. 2008). The pursuit area of the frontal eye field (FEF_sem_) could also contribute to anticipatory pursuit, because lesions in FEF could abolish the ipsilateral anticipatory initiation of pursuit (Macavoy et al. 1991). We hypothesize that expectation based on visual and/or motor history might be encoded in SEF and then combined with current sensory evidence in FEF (Darlington et al. 2018; Fukushima et al. 2013; Schall 2015). The source of visual motion history might come from MT and MST, but the roles of these areas in anticipatory pursuit remain to be tested (Kowler et al. 2019).

### Limitations and future directions

One limitation of the current study is that we were not able to analyze the temporal development of the expectation effect in perception in detail due to limited number of probe trials. Anticipatory pursuit and perception rely on history on different temporal scales (Maus et al. 2015). Attraction and repulsion perceptual bias seem to also operate on different time scales (Chopin and Mamassian 2012; Fritsche et al. 2020), and could occur with slight changes of parameters in the same paradigm (Kanai and Verstraten 2005). Therefore, understanding the temporal dependency and development of the effects of expectation would be crucial to understand the complicated interaction between attraction and repulsion biases.

We do not know how robust our results are with regard to parametric variations of the visual stimuli. For example, an attraction bias is mostly observed in studies using moving stimuli whose directions differed by about 60° or less (Fritsche et al. 2020) rather than 180° (as in our study). In addition, effects of expectation are often examined with weak motion stimuli, i.e., low contrast (Chalk et al. 2010) or low coherence (probe trials in our study), because the Bayesian integration hypothesis postulates that the effect of expectation would be larger on a stimulus with less reliable sensory signals. However, reducing coherence might introduce changes other than reducing contrast for RDK stimuli, such as inducing the perceptual phenomenon of motion transparency, in which two or more distinct surfaces are perceived as moving in different direction (Qian et al. 1994). Motion characteristics of noise dots in an RDK, together with their lifetime, affect the perception of global motion as well as pursuit quality (Pilly and Seitz 2009; Schütz et al. 2010). Future work is needed to elucidate the potential influence of the characteristics of sensory stimuli—ranging from simplified dots, blobs and RDKs to more complex naturalistic stimuli (Goettker et al. 2020)—on behavioral biases in perception and eye movements.

Finally, the perceptual repulsion bias observed in our and other studies does not match predictions of optimal Bayesian integration. Standard Bayesian inference would predict an attraction bias to the prior. Combining this prediction with the efficient coding hypothesis, whereby expectation modulates sensory likelihood, could account for the repulsion biases (“Anti-Bayesian” effects; Wei and Stocker 2015). In the future, this kind of modeling approach might help understand the complicated interaction between attraction and repulsion biases induced by experience-based expectation across different behavioral tasks.

## Acknowledgements

This work was supported by a University of British Columbia four-year fellowship to X.W., a Natural Sciences and Engineering Research Council of Canada Discovery Grant and Accelerator Supplement to M.S., and a CNRS - PICS grant “APPVIS” to A.M. Preliminary data from this project were presented at the 2019 Gordon Research Conference on Eye Movements, the 2019 Gordon Research Seminar on Eye Movements, the Society for Neuroscience annual meeting (Wu et al. 2019), and the Vision Sciences Society annual meeting (Wu et al. 2020). The authors thank members of the Spering lab for comments on an earlier draft of the article.

## References

Alais D, Leung J, Van der Burg E. Linear summation of repulsive and attractive serial dependencies: Orientation and motion dependencies sum in motion perception. J Neurosci 37: 4381–4390, 2017.

Bates D, Mächler M, Bolker BM, Walker SC. Fitting linear mixed-effects models using lme4. J Stat Softw 67: 1–48, 2015.

Behling S, Lisberger SG. Different mechanisms for modulation of the initiation and steady-state of smooth pursuit eye movements. J Neurophysiol 123: 1265–1276, 2020.

Born RT, Bradley DC. Structure and function of visual area MT. Annu Rev Neurosci 28: 157–189, 2005.

Brainard DH. The Psychophysics Toolbox. Spat Vis 10: 433–6, 1997.

Braun DI, Pracejus L, Gegenfurtner KR. Motion aftereffect elicits smooth pursuit eye movements. J Vis 6: 671–684, 2006.

Chalk M, Seitz AR, Series P. Rapidly learned stimulus expectations alter perception of motion. J Vis 10: 1–17, 2010.

Chopin A, Mamassian P. Predictive Properties of Visual Adaptation. Curr Biol 22: 622–626, 2012.

Cicchini GM, Mikellidou K, Burr DC. The functional role of serial dependence. Proc R Soc B Biol Sci 285: 1–8, 2018.

Damasse J-B, Perrinet LU, Madelain L, Montagnini A. Reinforcement effects in anticipatory smooth eye movements. J Vis 18: 1–18, 2018.

Darlington TR, Beck JM, Lisberger SG. Neural implementation of Bayesian inference in a sensorimotor behavior. Nat Neurosci 21: 1442–1451, 2018.

Darlington TR, Tokiyama S, Lisberger SG. Control of the strength of visual-motor transmission as the mechanism of rapid adaptation of priors for Bayesian inference in smooth pursuit eye movements. J Neurophysiol 118: 1173–1189, 2017.

Deravet N, Blohm G, de Xivry J-JO, Lefèvre P. Weighted integration of short-term memory and sensory signals in the oculomotor system. J Vis 18: 1–19, 2018.

Ferrera VP, Lisberger SG. Neuronal Responses in Visual Areas MT and MST During Smooth Pursuit Target Selection. J Neurophysiol 78: 1433–1446, 1997.

Fritsche M, Spaak E, de Lange FP. A Bayesian and efficient observer model explains concurrent attractive and repulsive history biases in visual perception. Elife 9, 2020.

Fukushima K, Fukushima J, Warabi T, Barnes GR. Cognitive processes involved in smooth pursuit eye movements: Behavioral evidence, neural substrate and clinical correlation. Front. Syst. Neurosci. 7Frontiers: 1–28, 2013.

Gegenfurtner KR. The Interaction Between Vision and Eye Movements. Perception 45: 1333–1357, 2016.

Goettker A, Agtzidis I, Braun DI, Dorr M, Gegenfurtner KR. From Gaussian blobs to naturalistic videos: Comparison of oculomotor behavior across different stimulus complexities. J Vis 20: 1–16, 2020.

Händel BH, Lutzenberger W, Thier P, Haarmeier T. Opposite Dependencies on Visual Motion Coherence in Human Area MT1 and Early Visual Cortex. Cereb Cortex 17: 1542–1549, 2007.

de Hemptinne C, Lefèvre P, Missal M. Neuronal bases of directional expectation and anticipatory pursuit. J Neurosci 28: 4298–4310, 2008.

Jarrett CB, Barnes G. Volitional selection of direction in the generation of anticipatory ocular smooth pursuit in humans. Neurosci Lett 312: 25–28, 2001.

Jarrett CB, Barnes G. Volitional scaling of anticipatory ocular pursuit velocity using precues. Cogn Brain Res 14: 383–388, 2002.

Jazayeri M, Movshon JA. A new perceptual illusion reveals mechanisms of sensory decoding. Nature 446: 912–915, 2007.

Kanai R, Verstraten FAJ. Perceptual manifestations of fast neural plasticity: Motion priming, rapid motion aftereffect and perceptual sensitization. Vision Res 45: 3109–3116, 2005.

Keck MJ, Palella TD, Pantle A. Motion aftereffect as a function of the contrast of sinusoidal gratings. Vision Res 16: 187–191, 1976.

Kleiner M, Brainard D, Pelli D., Ingling A, Murray R, Broussard C. What’s new in psychtoolbox-3. Perception 36: 1–16, 2007.

Kohn A, Movshon JA. Neuronal adaptation to visual motion in area MT of the macaque. Neuron 39: 681–691, 2003.

Kohn A, Movshon JA. Adaptation changes the direction tuning of macaque MT neurons. Nat Neurosci 7: 764–772, 2004.

Kok P, Brouwer GJ, van Gerven MAJ, de Lange FP. Prior expectations bias sensory representations in visual cortex. J Neurosci 33: 16275–16284, 2013.

Kowler E. Cognitive expectations, not habits, control anticipatory smooth oculomotor pursuit. Vision Res 29: 1049–1057, 1989.

Kowler E, Martins AJ, Pavel M. The effect of expectations on slow oculomotor control—IV. Anticipatory smooth eye movements depend on prior target motions. Vision Res 24: 197–210, 1984.

Kowler E, Rubinstein JF, Santos EM, Wang J. Predictive Smooth Pursuit Eye Movements. Annu Rev Vis Sci 5: 223–246, 2019.

Krauzlis RJ, Miles FA. Release of fixation for pursuit and saccades in humans: Evidence for shared inputs acting on different neural substrates. J Neurophysiol 76: 2822–2833, 1996.

Kreyenmeier P, Fooken J, Spering M. Context effects on smooth pursuit and manual interception of a disappearing target. J Neurophysiol 118: 404–415, 2017.

de Lange FP, Heilbron M, Kok P. How Do Expectations Shape Perception? Trends Cogn Sci 22: 764–779, 2018.

Lawrence MA. Easy analysis and visualization of factorial experiments [Online]. 2016.http://github.com/mike-lawrence/ez.

Macavoy MG, Gottlieb JP, Bruce CJ. Smooth-pursuit eye movement representation in the primate frontal eye field. Cereb Cortex 1: 95–102, 1991.

Madelain L, Krauzlis RJ. Pursuit of the ineffable: perceptual and motor reversals during the tracking of apparent motion. J Vis 3: 642–653, 2003.

Mather G, Pavan A, Campana G, Casco C. The motion aftereffect reloaded. Trends Cogn. Sci. 12: 481–487, 2008.

Maus GW, Potapchuk E, Watamaniuk SNJ, Heinen SJ. Different time scales of motion integration for anticipatory smooth pursuit and perceptual adaptation. J Vis 15: 1–13, 2015.

Meier K, Giaschi D. The maturation of global motion perception depends on the spatial and temporal offsets of the stimulus. Vision Res 95: 61–67, 2014.

Missal M, Heinen SJ. Supplementary Eye Fields Stimulation Facilitates Anticipatory Pursuit. J Neurophysiol 92: 1257–1262, 2004.

Montagnini A, Spering M, Masson GS. Predicting 2D Target Velocity Cannot Help 2D Motion Integration for Smooth Pursuit Initiation. J Neurophysiol 96: 3545–3550, 2006.

Pasture C, Montagnini A, Perrinet LU. Humans adapt their anticipatory eye movements to the volatility of visual motion properties. PLOS Comput Biol 16: e1007438, 2020.

Pelli DG. The VideoToolbox software for visual psychophysics: transforming numbers into movies. Spat Vis 10: 437–42, 1997.

Pilly PK, Seitz AR. What a difference a parameter makes: A psychophysical comparison of random dot motion algorithms. Vision Res 49: 1599–1612, 2009.

Prins N, Kingdom FAA. Applying the Model-Comparison Approach to Test Specific Research Hypotheses in Psychophysical Research Using the Palamedes Toolbox. Front Psychol 9: 1250, 2018.

Qian N, Andersen R, Adelson E. Transparent motion perception as detection of unbalanced motion signals. I. Psychophysics. J Neurosci 14: 7357–7366, 1994.

Santos EM, Gnang EK, Kowler E. Anticipatory smooth eye movements with random-dot kinematograms. J Vis 12: 1–1, 2012.

Santos EM, Kowler E. Anticipatory smooth pursuit eye movements evoked by probabilistic cues. J Vis 17: 1–16, 2017.

Schall JD. Visuomotor Functions in the Frontal Lobe. Annu Rev Vis Sci 1: 469–498, 2015.

Schütz AC, Braun DI, Gegenfurtner KR. Eye movements and perception: A selective review. J Vis 11: 1–30, 2011.

Schütz AC, Braun DI, Movshon JA, Gegenfurtner KR. Does the noise matter? Effects of different kinematogram types on smooth pursuit eye movements and perception. J Vis 10: 1–22, 2010.

Seriès P, Seitz AR. Learning what to expect (in visual perception). Front. Hum. Neurosci. 7Frontiers Media S. A.: 668, 2013.

Spering M, Gegenfurtner KR. Contrast and Assimilation in Motion Perception and Smooth Pursuit Eye Movements. J Neurophysiol 98: 1355–1363, 2007.

Spering M, Montagnini A. Do we track what we see? Common versus independent processing for motion perception and smooth pursuit eye movements: A review. Vision Res 51: 836–852, 2011.

Team RC. R: A language and environment for statistical computing [Online]. R Foundation for Statistical Computing2019.http://www.r-project.org/.

Thier P, Ilg UJ. The neural basis of smooth-pursuit eye movements. Curr Opin Neurobiol 15: 645–652, 2005.

Watamaniuk SNJ, Bal J, Heinen SJ. A Subconscious Interaction between Fixation and Anticipatory Pursuit. J Neurosci 37: 11424–11430, 2017.

Wei XX, Stocker AA. A Bayesian observer model constrained by efficient coding can explain “anti-Bayesian” percepts. Nat Neurosci 18: 1509–1517, 2015.

Wu X, Rothwell AC, Spering M, Montagnini A. Comparing dynamic effects of expectation on motion perception and pursuit eye movements. In: Program No. 226.12. 2019 Neuroscience Meeting Planner. Chicago, IL, United States: Society for Neuroscience, 2019.

Wu X, Rothwell AC, Spering M, Montagnini A. Opposite effects of expectation on motion perception and anticipatory pursuit eye movements. J Vis 20: 567, 2020.

Yang J, Lee J, Lisberger SG. The interaction of Bayesian priors and sensory data and its neural circuit implementation in visually guided movement. J Neurosci 32: 17632–17645, 2012.

